# Transcriptional cascades during fasting amplify gluconeogenesis and instigate a secondary wave of ketogenic gene transcription

**DOI:** 10.1101/2024.04.04.588039

**Authors:** Dana Goldberg, Nufar Buchshtab, Meital Charni-Natan, Ido Goldstein

## Abstract

**Background & Aims:** During fasting, bodily homeostasis is maintained due to hepatic production of glucose (gluconeogenesis) and ketone bodies (ketogenesis). The main hormones governing hepatic fuel production are glucagon and glucocorticoids that initiate transcriptional programs aimed at supporting gluconeogenesis and ketogenesis.

**Methods:** Using primary mouse hepatocytes as an ex vivo model, we employed transcriptomic analysis (RNA-seq), genome-wide profiling of enhancer dynamics (ChIP-seq), perturbation experiments (inhibitors, shRNA), hepatic glucose production measurements and computational analyses.

**Results:** We found that in addition to the known metabolic genes transcriptionally induced by glucagon and glucocorticoids, these hormones induce a set of genes encoding transcription factors (TFs) thereby initiating transcriptional cascades. Upon activation by glucocorticoids, the glucocorticoid receptor (GR) induced the genes encoding two TFs: CCAAT/enhancer-binding protein beta (C/EBPβ) and peroxisome proliferator-activated receptor alpha (PPARα). We found that C/EBPβ mainly serves as an amplifier of hormone-induced gene programs in hepatocytes. C/EBPβ augmented gluconeogenic gene expression and hepatic glucose production. Conversely, the GR-PPARα cascade initiated a secondary transcriptional wave of genes supporting ketogenesis. The cascade led to synergistic induction of ketogenic genes which is dependent on protein synthesis. Genome-wide analysis of enhancer dynamics revealed numerous enhancers activated by the GR-PPARα cascade. These enhancers were proximal to ketogenic genes, enriched for the PPARα response element and showed increased PPARα binding.

**Conclusion:** This study reveals abundant transcriptional cascades occurring during fasting. These cascades serve two separated purposes: the amplification of the primary gluconeogenic transcriptional program and the induction of a secondary gene program aimed at enhancing ketogenesis.

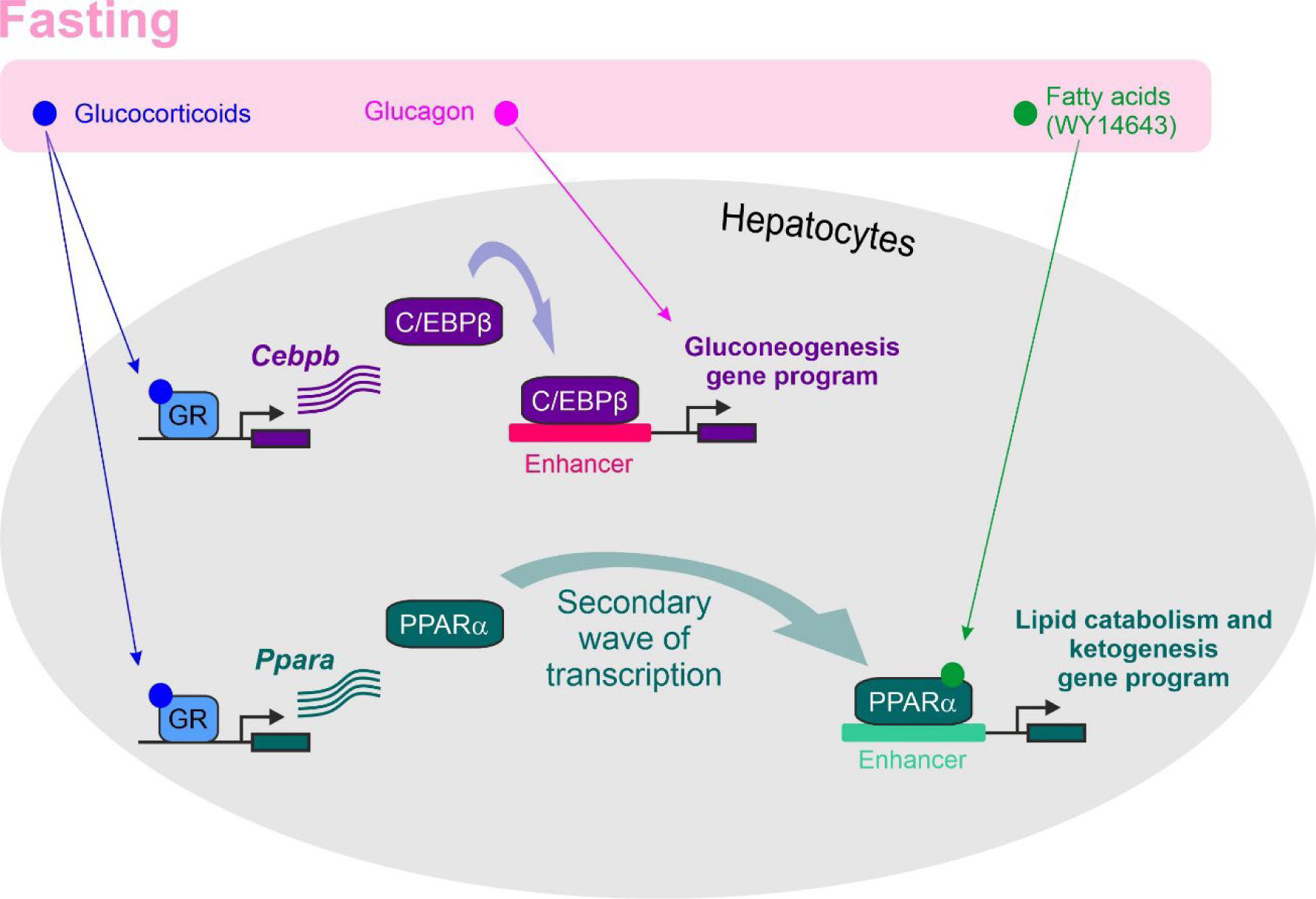

## Introduction

Mammals frequently undergo bouts of fasting during which their homeostasis is maintained by production of fuel from internal energy storages. The type of fuel produced depends on the length of fasting and is controlled by the liver. Fasting for a few hours elicits breaking down of glycogen and the subsequent release of glucose from hepatocytes to the blood, thereby supplying the needs of glucose-utilizing tissues such as the brain [1–3]. If fasting persists, glycogen stores are dwindled and glucose production is mostly achieved by gluconeogenesis, the de novo production of glucose from non-carbohydrate precursors such as amino acids. Because skeletal muscle proteins are a major source for gluconeogenic amino acids, over-reliance on gluconeogenesis puts muscle integrity at risk. Therefore, in prolonged fasting bouts, adipose tissue lipolysis is activated, triglycerides are broken down and free fatty acids travel to the liver where they are oxidized and provide precursors for ketogenesis. Ketone bodies produced in ketogenesis serve as efficient alternative fuel during fasting [3]. Thus, with the exhaustion of glycogen stores, gluconeogenesis provides glucose and ketogenesis bodies provide ketone bodies.

Fuel production during fasting is tightly regulated and orchestrated by endocrine and metabolic signals. Two principal hormones driving the fasting response and hepatic fuel production are glucagon [4] and glucocorticoids [5]. Glucagon is secreted from the pancreas and glucocorticoids (cortisol in humans and corticosterone in mice) are secreted from the adrenal gland following fasting. The secretion of both hormones prominently increases in prolonged fasting periods when glycogen is depleted and gluconeogenesis as well as ketogenesis assume the major role in providing fuel [6–10]. Glucagon and glucocorticoids work in a synergistic manner to activate gluconeogenesis and supply glucose during fasting [11–19].

Activation of fuel production during fasting is regulated at many levels, including increased transport of precursors from circulation into the liver, post-translational activation of enzymes as well as suppression of opposing metabolic pathways to prevent futile cycles. A principal regulatory node controlling fuel production is transcriptional regulation. Gluconeogenesis, fatty acid oxidation (FAO) and ketogenesis are all extensively regulated at the transcriptional level with hundreds of genes induced during fasting to support these metabolic pathways [20, 21]. The transcriptional programs initiated during fasting are controlled by a number of sequence-specific transcription factors (TFs) that are activated upon fasting. With regards to TFs, the term ‘activated’ is used to describe a TF that is bound to its DNA response element (motif) and promotes gene transcription [22]. Response elements reside within longer cis-regulatory elements termed enhancers. Enhancers can be adjacent to gene core promoters (and are therefore often termed promoter-proximal regions) or distal from them. Both distal and proximal enhancers serve the same purpose – they harbor clusters of response elements that, when bound by an activated TF, promote gene transcription [23, 24]. This is achieved by an intricate set of events initiated upon TF activation: TFs recruit co-activator proteins, histone-modifying enzymes and chromatin remodelers, leading to de-compaction chromatin in the enhancer area and an increase in certain histone acetyl marks such as H3K27 acetylation (H3K27ac) [25]. These events are broadly termed ‘enhancer activation’ and they eventually lead to increased rate of gene transcription by RNA polymerase II [22].

There are many manners by which TFs are activated during fasting. Glucagon elicits a signal transduction pathway leading to the phosphorylation of a TF termed cAMP response element-binding protein (CREB) [26]. CREB phosphorylation activates it, leading to the induction of gluconeogenic genes as well as amino acid catabolism genes providing precursors for gluconeogenesis [19, 26, 27]. Another TF, the glucocorticoid receptor (GR), is activated following its binding to glucocorticoids and the subsequent entry into the nucleus [28]. Hepatic GR supports gluconeogenesis during fasting by inducing pro-gluconeogenic genes [5, 20]. Both CREB and GR become active during fasting due to the increase in glucagon and glucocorticoids. A third TF activated by a fasting signal is peroxisome proliferator activated receptor alpha (PPARα), a ligand-activated TF. Fatty acids serve as a ligand to PPARα and they directly bind and activate it. Fatty acids influx into hepatocytes is rampant during fasting due to adipose tissue lipolysis and the subsequent release of fatty acids [29, 30]. In turn, PPARα induces hundreds of genes responsible for various aspects of lipid catabolism, FAO, and ketogenesis [31]. In fact, PPARα is such a central regulator of these pathways that its deletion impairs the ability of mice to withstand prolonged fasting periods [29].

While TFs can be activated by phosphorylation, nuclear localization and ligands; some TFs are constitutively active. This means that upon their translation, these TFs promote gene transcription constitutively. Therefore, regulating the expression of these TFs is key to controlling their activity. CCAAT enhancer binding protein beta (C/EBPβ) is a constitutively active TF which regulates key hepatic features [32, 33]. With regards to the fasting response, C/EBPβ and its close family member, C/EBPα, promote gluconeogenesis [34–39] and the malate-aspartate shuttle [40] which is needed to sustain gluconeogenic flux [41].

Upon activation, a TF induces a set of genes often termed ‘primary target genes’. Many of the genes transcriptionally-regulated in biological processes encode enzymes, transporters, secreted factors etc. A special group of genes are those encoding TFs. Induction of these genes instigates a ‘transcriptional cascade’ in which the induced TF-encoding gene is translated and upon its own activation, a secondary wave of gene transcription ensues. Thus, the first TF indirectly leads to the induction of ‘secondary target genes’ in a manner dependent on the synthesis and activation of the second TF. Transcriptional cascades play important roles in development where they control cell differentiation [42–44]. Fully-differentiated cells also partake in transcriptional cascades in order to optimally respond to environmental changes such as stress, endocrine signals and nutritional variations [18, 27, 45–47], although transcriptional cascades in differentiated cells are considerably less studied and are usually not considered principal nodes of gene regulation.

Here, we found that during fasting, the two principal fasting hormones glucagon and glucocorticoids induce a slew of TF-encoding genes, leading to transcriptional cascades. An in-depth investigation of two induced TFs reveals two roles for transcriptional cascades: the amplification of the primary gluconeogenic transcriptional program and the induction of a secondary gene program aimed at enhancing FAO and ketogenesis.

## Results

### Fasting hormones induce TF-encoding genes with central roles in the fasting response

The hepatic transcriptional regulation imposed by glucagon and glucocorticoids is centered around gluconeogenesis and its supporting pathways. To explore all aspects of this hormone-controlled gene program, we utilized our previously-published transcriptome profiling experiment in which we treated primary mouse hepatocytes with glucagon, corticosterone (the principal glucocorticoid in mice), or both hormones together in a dual treatment for 3 h. Of note, the durations of hormone treatments used throughout the manuscript aim to reflect different stages of prolonged fasting rather than short-term fasting. In short-term fasting glycogen breakdown provides most fuel. Glucagon and corticosterone only increase after ∼16 h of fasting [6–10]. Therefore, hormone treatments of 2-8 h used in this ex vivo study reflect mid-term to prolonged fasting (16-24 h in mice). Our previous RNA-seq profiling exposed widespread regulation of gene expression by both glucagon and corticosterone as well as a synergistic induction of genes in the dual treatment [17]. Here, to further explore the gene programs governed by glucagon and corticosterone, we searched for Gene Ontology (GO) terms enriched among induced genes. As expected, most enriched GO terms were related to metabolism and particularly to fasting-related biochemical pathways such as gluconeogenesis. This aligns with the well-established role of glucagon and corticosterone in augmenting gluconeogenesis. A prominent deviation from metabolic terms among hormone-induced genes was the high enrichment of GO terms relating to transcriptional regulation. In fact, 24% of the top enriched terms in glucagon-induced genes were related to transcriptional regulation with a similar fraction (21%) found in corticosterone-induced genes (Table S1). This suggests that the transcriptional programs controlled by glucagon and corticosterone are focused not only on directly regulating metabolic genes but also on inducing genes whose encoded proteins themselves regulate gene transcription.

Inspection of genes belonging to transcriptional regulation GO terms revealed that most of them encode sequence-specific TFs (i.e., TFs directly binding response elements in the genome) which we briefly term here TFs. To quantify the number of TFs induced by fasting hormones, we overlapped our lists of induced genes with a list of all mouse TF-encoding genes (n = 1,636) [48]. As pointed by the GO terms enrichment analysis, we found a clear enrichment of TF-encoding genes among hormone-induced genes with 14% of glucagon-induced genes encoding TFs and a similar fraction in corticosterone- and dual-induced genes (13% and 12%, respectively). A similar pattern was observed in genes repressed by these treatments as 11-19% of repressed genes encoded TFs (Table S1). To examine the relevance of this gene regulation pattern to in vivo fasting, we compared the TF-encoding genes induced in our dataset to genes induced in the liver following 24 h of fasting compared to a fed control [18]. We found noteworthy overlap between the two datasets as a total of 29 TFs were induced both during fasting and following at least one fasting hormone (Table S1). Some of the induced TFs play a documented role in the fasting response (e.g., FoxO1 [49–51] and Klf15 [52–54]). Similarly to the overlap in fasting-induced genes, we found that 13 TF-encoding genes repressed by fasting hormones are also repressed by fasting in mouse liver (Table S1), some of which were previously reported to inhibit fasting-related processes in the liver (e.g., FoxQ1 [55]). These findings show that fasting hormones profoundly regulate the levels of hepatic TFs and suggest that in addition to directly inducing metabolic pathways, these hormones also induce TFs that in turn may contribute to the fasting response. Examples for TFs induced both during fasting and following at least one fasting hormone are shown in Fig. 1A.

**Figure 1:**
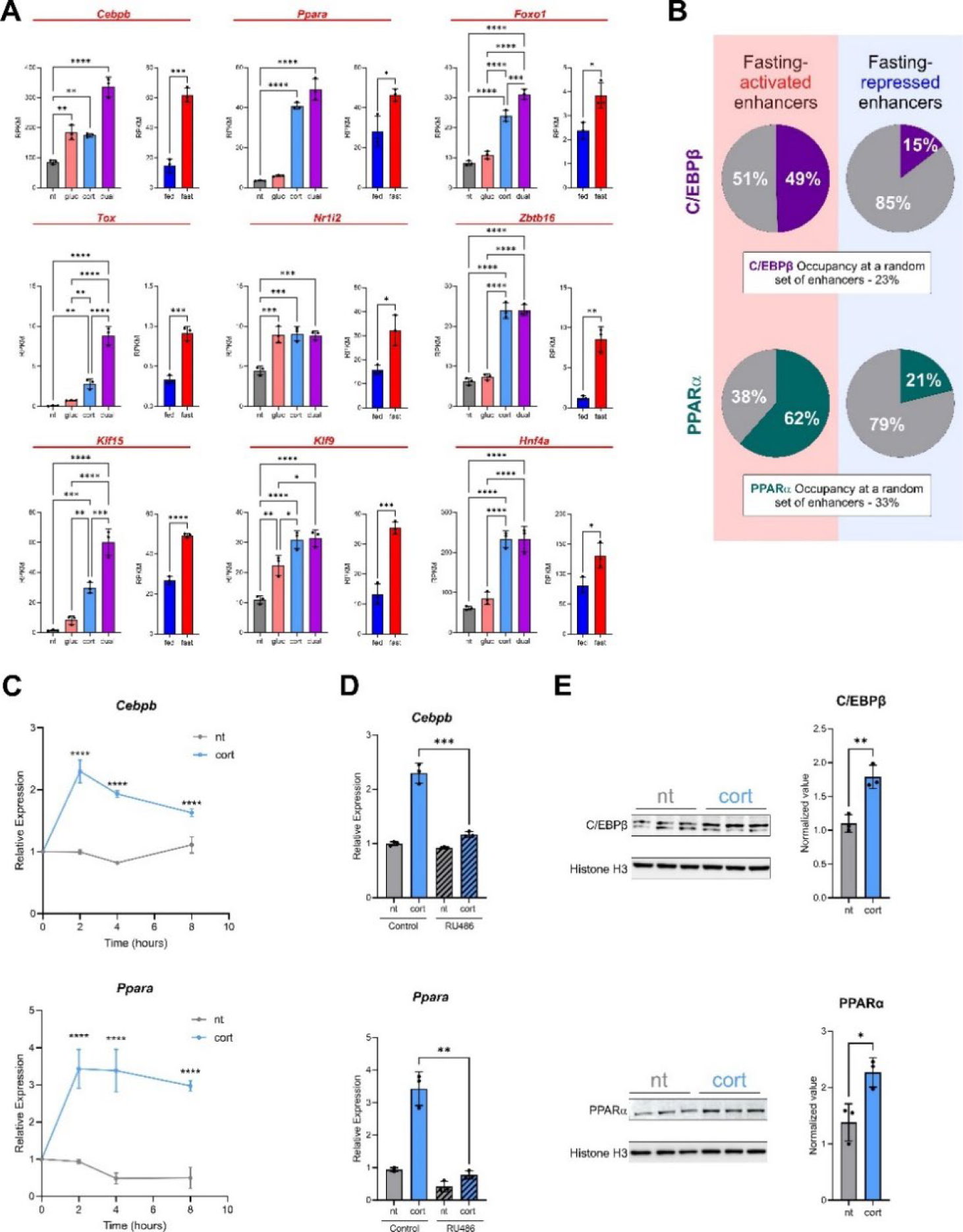
C/EBPβ and PPARα are rapidly and directly induced by the corticosterone-GR axis. **A.** The gene levels of selected hormone-induced, fasting-induced TF-encoding genes following a 3 h treatment of corticosterone (cort) or glucagon (gluc) in primary mouse hepatocytes or following 24 h of fasting in mouse liver. **B.** Overlap between fasting-regulated enhancers and TF binding sites shows preferential binding of both C/EBPβ and PPARα at fasting-activated enhancers as compared to either fasting-repressed enhancers or a random set of enhancers. **C.** Time course experiment shows that the induction of *Cebpb* and *Ppara* by corticosterone is immediate and occurs as early as 2 h following corticosterone treatment. Statistical tests were performed to compare between corticosterone-treated group and the non-treated group within each time point. **D.** Primary hepatocytes were treated with RU486 (a GR antagonist) prior to a 2 h treatment with corticosterone. The induction of *Cebpb* and *Ppara* by corticosterone is GR-dependent as it is negated in the presence of RU486. **E.** The protein levels of C/EBPβ and PPARα were assessed by western blot, showing an increase in protein levels 8 h following corticosterone treatment. - *P≤0.05, **P≤0.01, ***P≤0.001, ****P≤0.0001 by ordinary one-way ANOVA followed by Holm-Sidak post hoc analysis (A, B) or a two-tailed, unpaired t-test (C, E). RPKM: reads per kilobase per million reads.

Two notable TFs induced by corticosterone and during fasting were C/EBPβ (encoded by *Cebpb*) and PPARα (encoded by *Ppara*; Fig. 1A). This finding is in agreement with previous reports [56–59]. Glucagon also induced *Cebpb* (with an additive effect seen with corticosterone in the dual treatment) while *Ppara* levels were unresponsive to glucagon. Both C/EBPβ and PPARα were shown in various reports to induce fasting-related gene expression [31, 34–38] and were among the four TFs found to be associated with widespread changes in hepatic chromatin accessibility and enhancer activation following fasting [18]. To independently show the genome-wide association of C/EBPβ and PPARα with fasting-activated enhancers, we overlapped the genome-wide binding events of either C/EBPβ [18] or PPARα [60] with a set of fasting-activated enhancers (determined by a fasting-dependent increase in chromatin accessibility [19]). We found that 49% of fasting-activated enhancers are occupied by C/EBPβ and 62% are occupied by PPARα. This is in contrast to much lower occupancy rates in fasting-repressed enhancers or randomly-selected enhancers (Fig. 1B). In accordance with occupancy events along fasting-activated enhancers, the average occupancy levels measured by ChIP-seq signal strength of both TFs were also significantly higher at fasting-activated enhancers as compared to either fasting-repressed enhancers or randomly-selected enhancers (Fig. S1A). The same pattern was observed when instead of chromatin accessibility, we profiled the active enhancer mark H3K27ac (Fig. S1A). Therefore, in line with their documented role in the fasting response, C/EBPβ and PPARα preferentially bind fasting-activated enhancers in the liver.

Similarly to C/EBPβ and PPARα, two other TFs were previously found to be associated with fasting-activated enhancers: CREB (activated by glucagon) and GR (activated by corticosterone) [18]. When considered together, it is plausible that among the four key fasting-related TFs, CREB and GR are rapidly activated by glucagon and corticosterone and in addition to inducing gluconeogenic genes, they induce the two other key TFs -C/EBPβ and PPARα. In that way, fasting hormones elicit two waves of transcriptional regulation by eliciting transcriptional cascades. To explore this possibility we focused on the hormone-dependent induction of C/EBPβ and PPARα and its downstream effects. To untangle the copious effects of fasting, we used an ex vivo model where primary mouse hepatocytes were treated with fasting hormones. Because corticosterone induces both *Cebpb* and *Ppara* and therefore initiates both cascades we continued our investigation into corticosterone-dependent TF induction.

To assess the kinetics of TF induction, we performed a time-course experiment and found that the induction of *Cebpb* and *Ppara* peaks at 2 h following corticosterone treatment and wanes at later time points (Fig. 1C), suggesting a rapid and direct induction by GR following its activation by corticosterone. As expected, inhibition of GR with an antagonist (RU486) abolished the induction of *Cebpb* and *Ppara* by corticosterone (Fig. 1D). In addition, analysis of GR chromatin immunoprecipitation sequencing (ChIP-seq) data showed binding of GR in the vicinity of *Cebpb* and *Ppara* following corticosterone treatment in primary hepatocytes and in mouse liver following fasting with these sites also containing the GR response element (GRE; Fig. S1B). The gene induction is followed by an increase in the protein levels of both C/EBPβ and PPARα, evident 8 h following corticosterone treatment (Fig. 1E). Taken together, these findings demonstrate that the immediate response to corticosterone during fasting is characterized by extensive induction of TF-encoding genes, among them are C/EBPβ and PPARα that are rapidly and directly induced by GR.

### C/EBPβ amplifies the hormone-controlled gluconeogenic gene program

C/EBPβ controls various processes in hepatocytes with one of the most prominent pathways being promotion of gluconeogenesis through inducing gluconeogenic genes. Glucagon and corticosterone also promote gluconeogenic gene expression by activating CREB and GR, respectively. Due to the well-established role of C/EBPβ, glucagon and corticosterone in promoting gluconeogenic gene expression, we hypothesized that the induction of C/EBPβ serves to support and strengthen the primary gluconeogenic response initiated by glucagon and corticosterone. To explore this possibility, we aimed to alter C/EBPβ activity and examine its effect on gene expression. C/EBPβ is constitutively active and therefore it is mostly regulated at the level of expression. Therefore, we chose to alter C/EBPβ activity by reducing its expression levels using adenovirally-infected short hairpin RNA targeting C/EBPβ (Ad-shCebpb) which led to a 10-fold reduction in *Cebpb* levels (Fig. 2A). To study the effect of C/EBPβ on hormone-regulated gene expression, we treated primary hepatocytes with glucagon, corticosterone or a dual treatment in the presence of Ad-shCebpb. To capture the effect of induced C/EBPβ and not only basal C/EBPβ levels, we treated hepatocytes with hormones for 8 h, which is sufficient to elevate C/EBPβ protein levels (Fig. 1E). We profiled the transcriptome of primary hepatocytes under these conditions via RNA-seq and performed a set of pairwise comparisons to examine the effect of C/EBPβ on gene expression: we quantified differential expression between the Ad-Control and Ad-shCebpb hepatocytes in all conditions (non-treated, glucagon, corticosterone and dual). We found significant C/EBPβ-dependent effects on gene expression with a total of 1,084 genes regulated by C/EBPβ in at least one condition (Fig. 2B, Table S2). To differentiate between hormone-induced genes and C/EBPβ-regulated genes we use the following terminology: genes whose levels increase in the Ad-Control condition as compared to Ad-shCebpb were termed C/EBPβ-upregulated genes; genes whose levels decrease in the Ad-Control condition as compared to Ad-shCebpb were termed C/EBPβ-downregulated genes.

**Figure 2:**
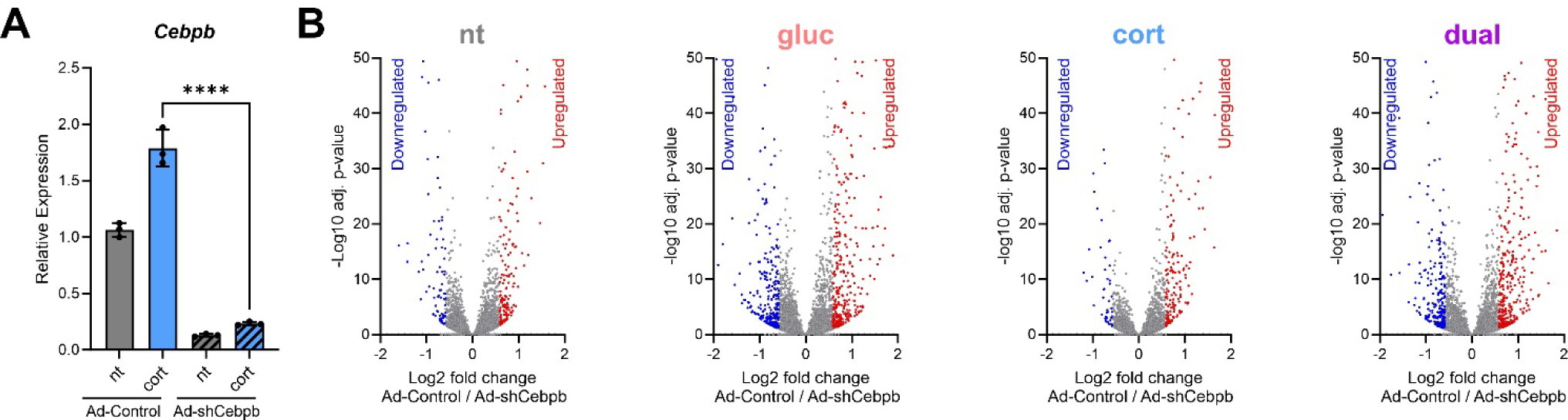
C/EBPβ has a widespread effect on hepatic gene expression. **A.** Adenovirally-infected shRNA targeting *Cebpb* (Ad-shCebpb) efficiently reduces its levels. **B.** Primary mouse hepatocytes were infected with Ad-shCebpb or a control (Ad-Control). The following day, cells were treated for 8 h with hormones as indicated and profiled by RNA-seq. Differential expression analysis between Ad-Control and Ad-shCebpb revealed a widespread effect of C/EBPβ on hepatocyte gene expression. - *P≤0.05, **P≤0.01, ***P≤0.001, ****P≤0.0001 by a two-tailed, unpaired t-test.

To evaluate the effect of C/EBPβ across different conditions, we performed gene clustering on all C/EBPβ-regulated genes. This revealed 8 clusters representing different gene expression patterns (Fig. 3A). Notably, only a minority of genes were regulated by C/EBPβ regardless of hormone treatment (Clusters 4, 6). The vast majority of C/EBPβ-regulated genes were concomitantly regulated by fasting hormones. The prominent tendency of C/EBPβ was to support and reinforce the regulation imposed by glucagon and corticosterone. Indeed, in Clusters 1, 2, 3, 5 and 8 the presence of C/EBPβ amplified the effect of hormone-dependent gene regulation (be it induction or repression). Only in cluster 7, C/EBPβ antagonized hormone-dependent gene induction. In other words, the role of C/EBPβ in hepatocytes is primarily focused on strengthening gene regulation imposed by fasting hormones (an example gene from each cluster is shown in Fig. 3B).

**Figure 3:**
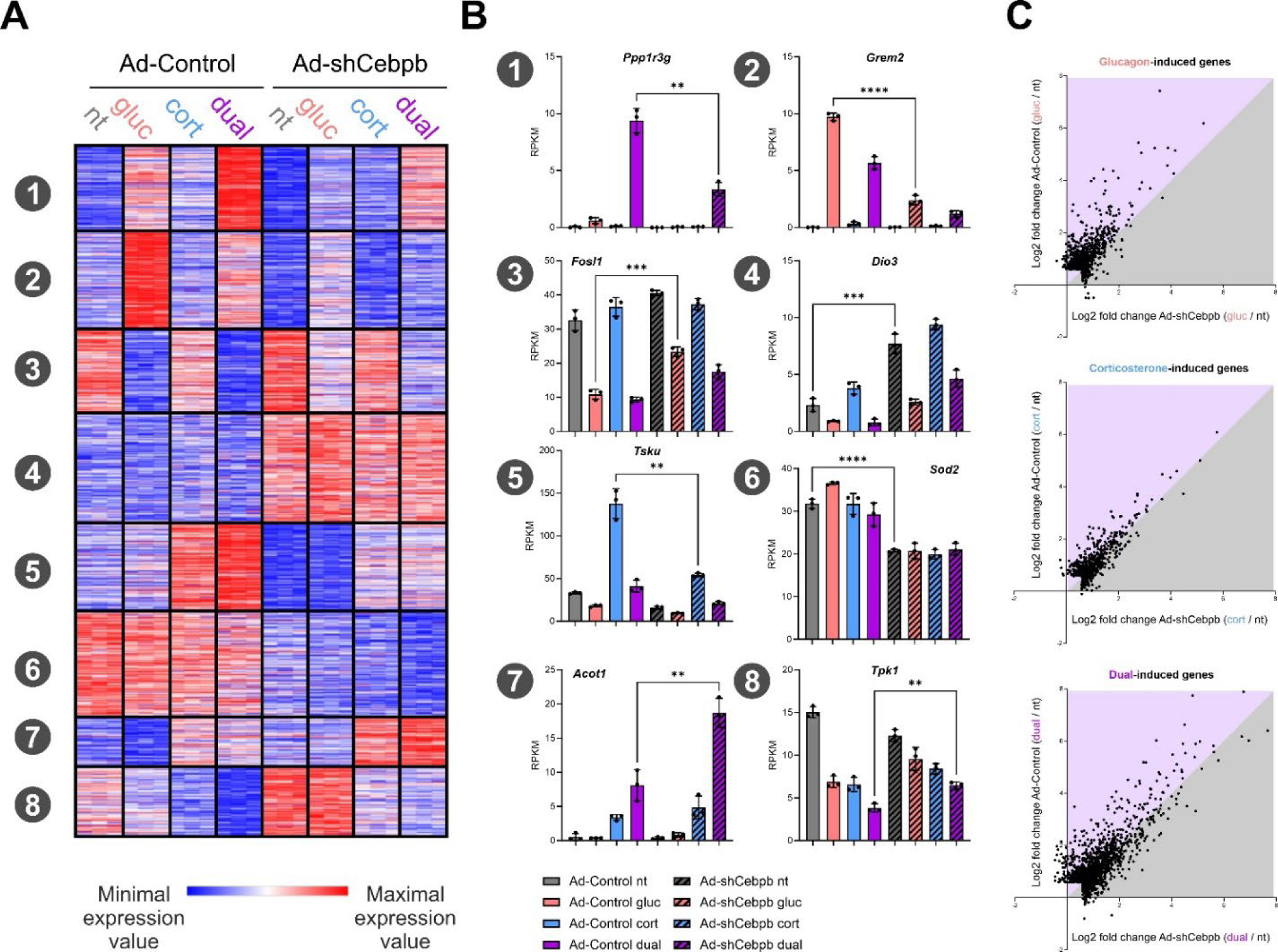
A principal role of C/EBPβ in hepatocytes is to amplify hormone-dependent gene regulation. **A.** k-means clustering of C/EBPβ-regulated genes (n = 1,084; k = 8) shows various gene expression patterns: *(1)* Dual-induced genes whose induction is strengthened by C/EBPβ *(2)* Glucagon-induced genes whose induction is strengthened by C/EBPβ *(3)* Glucagon-repressed genes whose repression is strengthened by C/EBPβ *(4)* C/EBPβ-downregulated genes *(5)* Corticosterone-induced genes whose induction is strengthened by C/EBPβ *(6)* C/EBPβ-upregulated genes *(7)* Dual-induced whose induction is antagonized by C/EBPβ *(8)* Dual-repressed genes whose repression is strengthened by C/EBPβ Blue: minimum expression value of gene. Red: maximum expression value of gene (minimum and maximum values of each gene are set independently to other genes). **B.** The expression of an example gene from each cluster is shown. **C.** The fold change values for all hormone-induced genes are shown (i.e., genes induced by glucagon, corticosterone or dual treatments compared to the non-treated control). The fold change values in Ad-Control cells are plotted along the x axis and the fold change values in Ad-shCebpb cells are plotted along the y axis. Comparing the induction capacity of genes in the presence or absence of C/EBPβ reveals a supporting role for C/EBPβ in hormone-dependent gene induction with many genes showing higher induction in the presence C/EBPβ (lilac-shaded area). The most prominent effect is observed in glucagon-induced genes. - *P≤0.05, **P≤0.01, ***P≤0.001, ****P≤0.0001 by a two-tailed, unpaired t-test. RPKM: reads per kilobase per million reads.

To evaluate on which hormone C/EBPβ exerts a stronger effect, we plotted the fold induction of all hormone-induced genes over the non-treated control. The fold induction in the presence of Ad-shCebpb (x axis) versus the fold induction in the presence of Ad-Control (y-axis) was plotted. Thus, a gene whose hormone-dependent induction is augmented by C/EBPβ (i.e. a gene with higher fold induction in the presence of Ad-Control compared to Ad-shCebpb) would appear above the 45° diagonal. The plots reveal that the highest augmenting effect of C/EBPβ is observed on glucagon-induced genes with a more modest effect on corticosterone-induced genes (Fig. 3C). This effect was also evident in the total number of glucagon-induced genes that were concomitantly upregulated by C/EBPβ (n = 203) compared with the total number of corticosterone-induced genes concomitantly upregulated by C/EBPβ (n = 77; Table S3).

Next, we aimed to explore the biological roles of C/EBPβ-amplified, hormone-induced genes. GO terms enrichment analysis of C/EBPβ-amplified corticosterone-induced genes revealed a strong enrichment of genes encoding acute phase reactants, including the gene encoding C-reactive protein (Fig. 4A; Table S3). Acute phase reactants are produced and secreted by hepatocytes during infection and are critical in restraining infection and preventing tissue damage [61–63]. Thus, production of acute phase reactants is one of the most important liver functions [61]. A role for C/EBPβ and GR in the acute phase response is long-known [62, 64] and our findings link the effects of these two TFs, suggesting they cooperate to induce acute phase genes.

**Figure 4:**
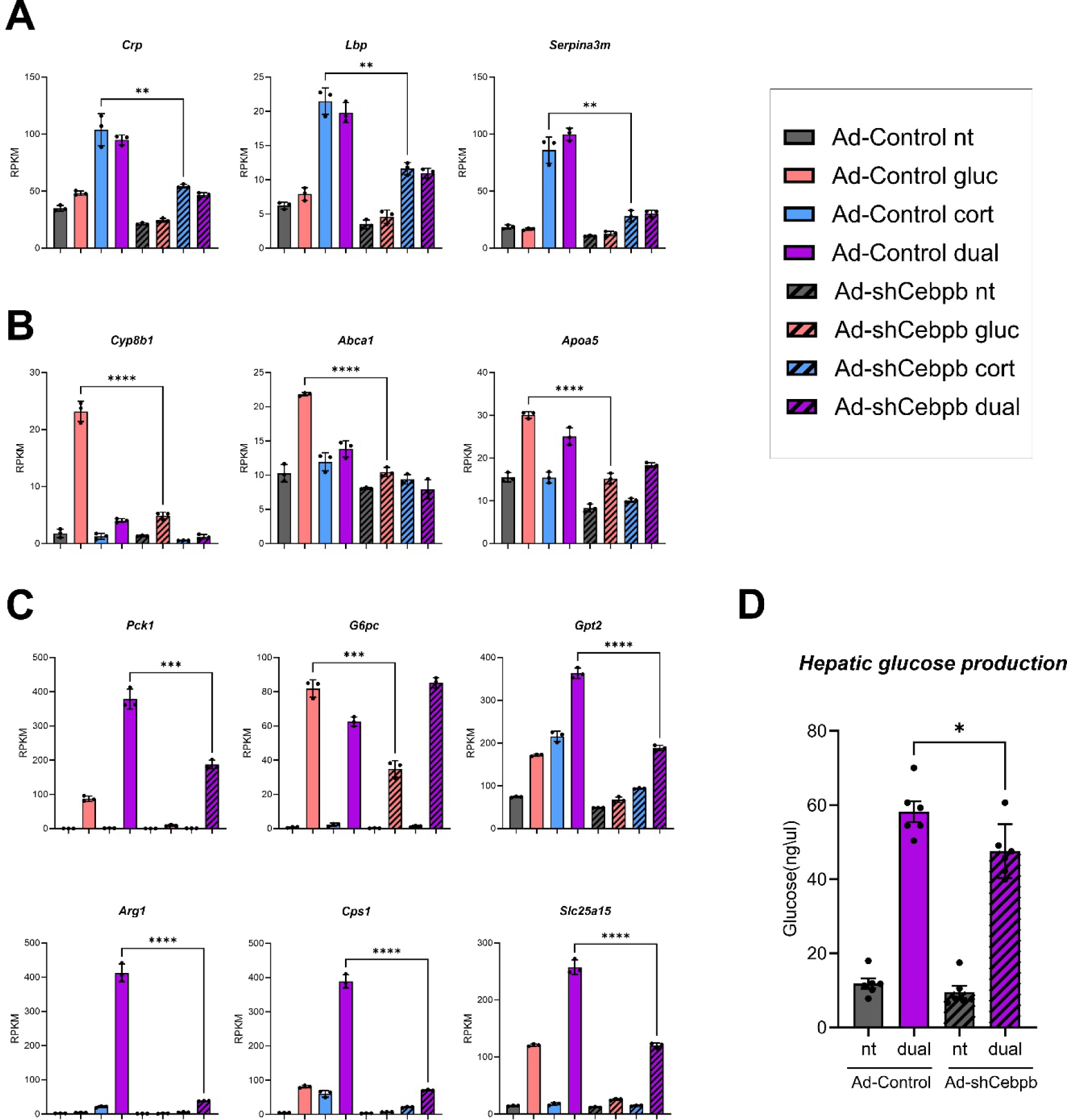
C/EBPβ augments gluconeogenic gene expression and hepatic glucose production. **A.** Quantification of acute-phase response genes shows C/EBPβ strengthens corticosterone-dependent gene induction. **B.** Quantification of cholesterol/bile acid metabolism genes shows C/EBPβ strengthens glucagon-dependent gene induction. **C.** Quantification of gluconeogenic genes shows C/EBPβ strengthens glucagon-corticosterone gene induction. **D.** Glucose produced from mouse hepatocytes following a dual treatment (glucagon and corticosterone) was measured, showing cells produce more glucose in the presence of C/EBPβ. -Examples for more genes appear in Fig. S2. - *P≤0.05, **P≤0.01, ***P≤0.001, ****P≤0.0001 by a two-tailed, unpaired t-test. RPKM: reads per kilobase per million reads.

GO terms enrichment analysis of C/EBPβ-amplified glucagon-induced genes revealed a strong enrichment of terms related to bile acid metabolism and cholesterol transport. These include genes encoding key factors in bile acid metabolism (*Cyp8b1*, *Abcg5*, *Abcg8*) as well as lipoprotein metabolism (*Abca1*, *Apoa5*, *Abca8b*; Fig. 4B; Fig. S2A; Table S3). Fasting was reported to affect bile acid metabolism and transport [65, 66] and we show here that cooperation between glucagon and C/EBPβ drives the induction of related genes.

In addition to bile acid metabolism, C/EBPβ-amplified glucagon-induced genes were enriched for gluconeogenesis and pathways serving to provide gluconeogenic precursors (urea cycle and amino acid catabolism; Table S3). Many gluconeogenic and amino acid catabolism genes are synergistically-induced by glucagon and corticosterone [19]. Notably, the synergistic induction of four urea cycle genes (*Cps1, Arg1, Asl, Slc25a15*) was strongly inhibited when C/EBPβ levels were reduced as was the induction of gluconeogenic genes (*Pck1, G6pc*) and amino acid catabolism genes providing gluconeogenic precursors (*Gpt2, Tat, Mat1a*; Fig. 4C, Fig. S2B). Examining C/EBPβ occupancy at the genes’ loci showed increased C/EBPβ binding following fasting in mouse liver, pointing to a direct role of C/EBPβ in inducing these genes in liver (Fig. S2C).

Given the strong support C/EBPβ exerts on transcriptional programs driving gluconeogenesis and amino acid catabolism, we hypothesized it also augments the capacity of hepatocytes to produce glucose from amino acids. Thus, we measured the effect of C/EBPβ on glucose produced by hepatocytes from an amino acid precursor (glutamine). We treated primary hepatocytes with glucagon and corticosterone to maximally augment glucose production [19]. We found that the gluconeogenic capacity of Ad-shCebpb hepatocytes is impaired compared to that of hepatocytes with intact levels of C/EBPβ (Fig. 4D). Collectively, these findings show that a principal role of C/EBPβ in hepatocytes is to amplify the transcriptional response to fasting hormones and in particular augment gluconeogenic gene expression and subsequent glucose production.

### Mutual activation of PPARα and GR leads to synergistic induction of ketogenic genes

In contrast to C/EBPβ, the major and most heavily-documented role of PPARα during fasting is to activate lipid catabolism, FAO, and ketogenesis. Also unlike C/EBPβ, PPARα is a ligand-activated TF and is only fully functional following binding to its ligand. Therefore, to explore the link between the corticosterone-dependent induction of *Ppara* and the ensuing activity of PPARα we treated hepatocytes with either of the following treatments: corticosterone alone, Wy-14643 treatment (a PPARα-specific ligand [67], abbreviated here to Wy) or a dual treatment of both corticosterone and Wy. We treated hepatocytes for 8 h, which is sufficient to elevate PPARα protein levels (Fig. 1E). This experimental design allows to tease out the effects of GR alone (corticosterone treatment), activated PPARα at basal expression levels prior to *Ppara* induction (Wy treatment) or the combined effects of activated GR and activated PPARα following *Ppara* induction and the subsequent accumulation of PPARα (dual treatment). To measure changes in gene expression, we performed RNA-seq followed by differential gene expression analysis. Corticosterone treatment led to prominent gene induction with 1,046 genes induced. Much fewer genes were induced by Wy (n = 132). To test our hypothesis that corticosterone increases PPARα-dependent gene induction, we measured gene induction in the dual treatment. From a total of 1,134 dual-induced genes, most were also induced in either of the single treatments, but 305 genes were only induced in the dual treatment (Fig. 5A; Table S4). This suggests a synergistic cooperation between the two treatments. To test for synergistic gene induction, we used previously-formulated cutoffs [17] based on the definition of synergy: the effect of the dual treatment is higher than the summed effect of both single treatments. Thus, in order for a gene to be determined as synergistically-induced, it has to fulfill these four conditions: the gene is induced in the dual treatment compared to the: (a) non-treated control, (b) the corticosterone single treatment and (c) the Wy single treatment. (d) In addition, for a gene to be defined as synergistic, its increase in expression in the dual treatment must be at least 1.3 higher than the *sum* of increases in the two single treatments combined (Fig. 5B). Using these strict cutoffs that combine several statistical cutoffs and a fold change threshold ensures that only bona-fide synergistically-induced genes are included and not additively-induced genes. This analysis revealed 80 genes that are synergistically-induced by a combined treatment of corticosterone and Wy (Table S4). GO terms enrichment analysis revealed absolute dominance of lipid catabolism gene programs that are governed by PPARα (Table S4). This included key genes involved in FAO, ketogenesis and triglyceride catabolism (Fig. 5C). The synergistic induction is completely dependent on GR as it is abolished in the presence of a GR antagonist (Fig. 5D).

**Figure 5:**
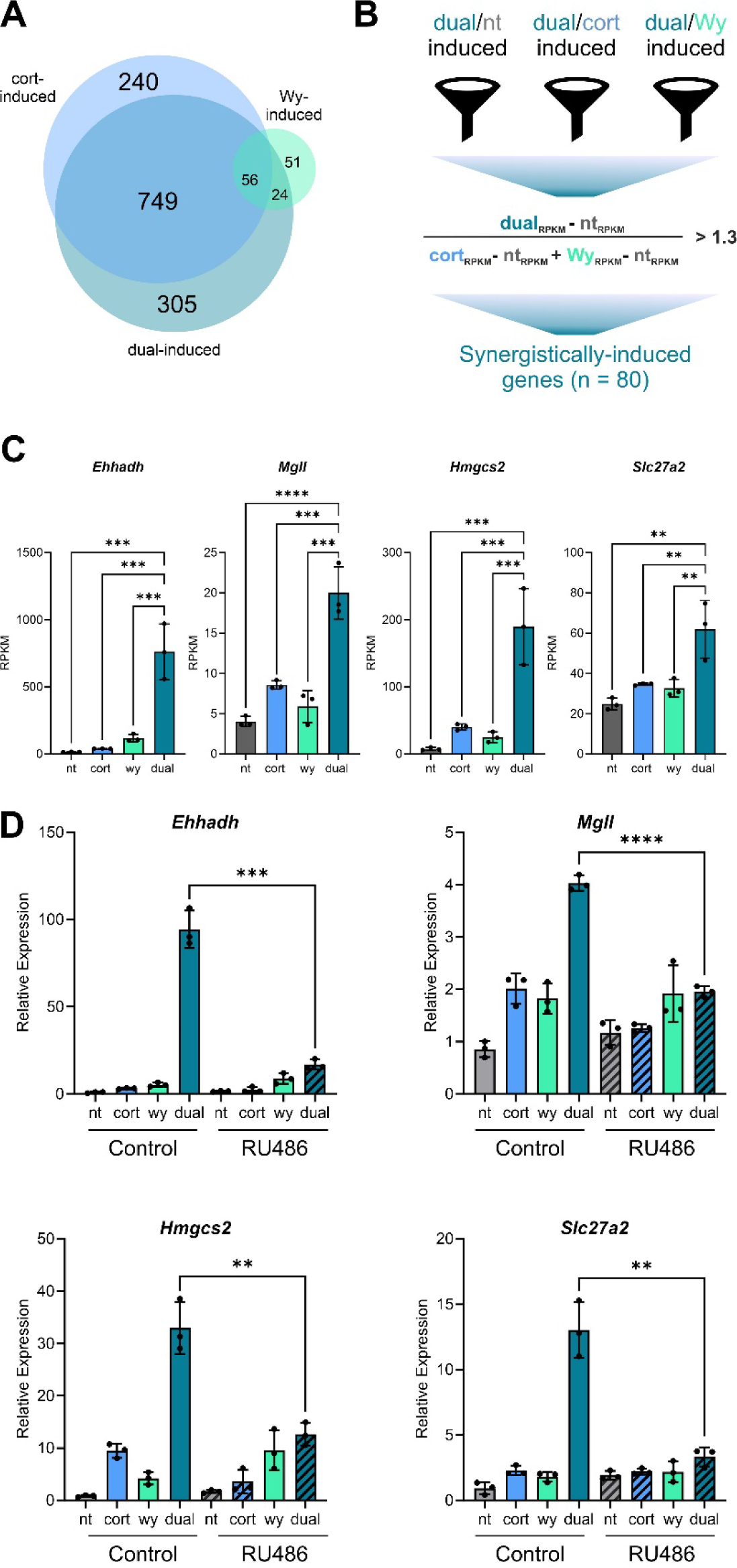
GR and PPARα synergistically-induce lipid catabolism and ketogenesis genes. **A.** Primary mouse hepatocytes were treated with corticosterone, Wy, or both in a dual treatment for 8 h. Transcriptome was profiled via RNA-seq and the set of induced genes in each condition was determined by comparing gene expression to the non-treated (nt) condition. The list of induced genes in each treatment were overlapped, showing 305 genes are only induced in the presence of both Wy and corticosterone. **B.** A synergistic relationship between Wy and corticosterone was defined with clear cutoffs: A synergistically-induced gene must meet 4 criteria: induction by the dual treatment compared to the non-treated control, compared to Wy and compared to corticosterone. Moreover, the gene expression increase in the dual treatment must be at least 1.3 higher than the sum of increased gene expression of the two single treatments. **C.** Examples for synergistically-induced genes are shown. **D.** Primary hepatocytes were treated with RU486 (a GR antagonist) prior to an 8 h treatment with corticosterone and Wy. Synergistic gene induction is dependent on GR as it is negated in the presence of RU486. - *P≤0.05, **P≤0.01, ***P≤0.001, ****P≤0.0001 by ordinary one-way ANOVA followed by Holm-Sidak post hoc analysis. RPKM: reads per kilobase per million reads.

### The GR-PPARα cascade leads to widespread enhancer activation

The enrichment PPARα-related GO terms together with the corticosterone-dependent increase in PPARα levels led us to the following hypothesis: synergistic gene induction is brought about by a corticosterone-dependent increase in PPARα protein levels, leading to an augmented cellular response to Wy thanks to the high levels of PPARα able to bind it. To test this hypothesis, we took several complementary approaches detailed below, including genome-wide enhancer profiling, perturbation experiments and a ‘PPARα replacement’ strategy.

First, we aimed to examine how the dual treatment of corticosterone and Wy differentially affects enhancer activation compared to the single treatments. We performed ChIP-seq for H3K27ac, a well-established marker for active enhancers. As expected, H3K27ac sites reside both in promoter-proximal and distal sites (Fig. S3A). H3K27ac signal in the loci of synergistically-induced genes was markedly increased in the dual treatment (Fig. 6A). To show this quantitatively across all synergistically-induced genes, we measured H3K27ac signal in the vicinity of all 80 synergistically-induced genes and found that neither Wy nor corticosterone were able to increase H3K27ac signal alone. Only the dual treatment caused H3K27ac to increase above the non-treated control, showing that the presence of both signals is necessary for activation of promoter-proximal regions of synergistic genes (Fig. 6B). Zooming out of promoter-proximal regions and examining enhancer activation on a genome-wide scale, we found a total of 1,971 enhancers were activated by corticosterone and only 35 enhancers were activated by Wy. The minimal level of enhancer activation by Wy aligns with minimal Wy-dependent gene induction. The dual treatment led to activation of 3,281 enhancers (Table S5). Some dual-activated enhancers were synergistically-activated (based on the same synergistic definitions made for gene induction, Table S5). As expected, the GR response-element (GRE) was the top enriched motif among corticosterone-activated enhancers and the PPARα response element (PPARE) was the top enriched motif in Wy-activated enhancers. In accordance, dual-activated enhancers were enriched with both motifs. A notable difference was found in synergistically-activated enhancers in which the GRE was absent and only the PPARE was enriched (Fig. 6C, Table S5). This suggests that synergistically-activated enhancers preferentially bind PPARα. While we and others were successful in performing ChIP-seq for PPARα in mouse liver, we were unable to obtain a high-quality ChIP-seq profile for PPARα in primary mouse hepatocytes. Therefore, we evaluated the binding of PPARα in synergistically-activated enhancers based on a previously-published, high-quality PPARα ChIP-seq experiment done in mouse liver in the fed and fasted states [60]. We found that PPARα binds synergistically-activated enhancers and that this binding significantly increases during fasting (Fig. 6A, S3B). Taken together, these observations show that lipid catabolism-related enhancers are only activated following a dual corticosterone and Wy treatment. Also, these enhancers are enriched with PPARE and are bound by PPARα during fasting in mouse liver.

**Figure 6:**
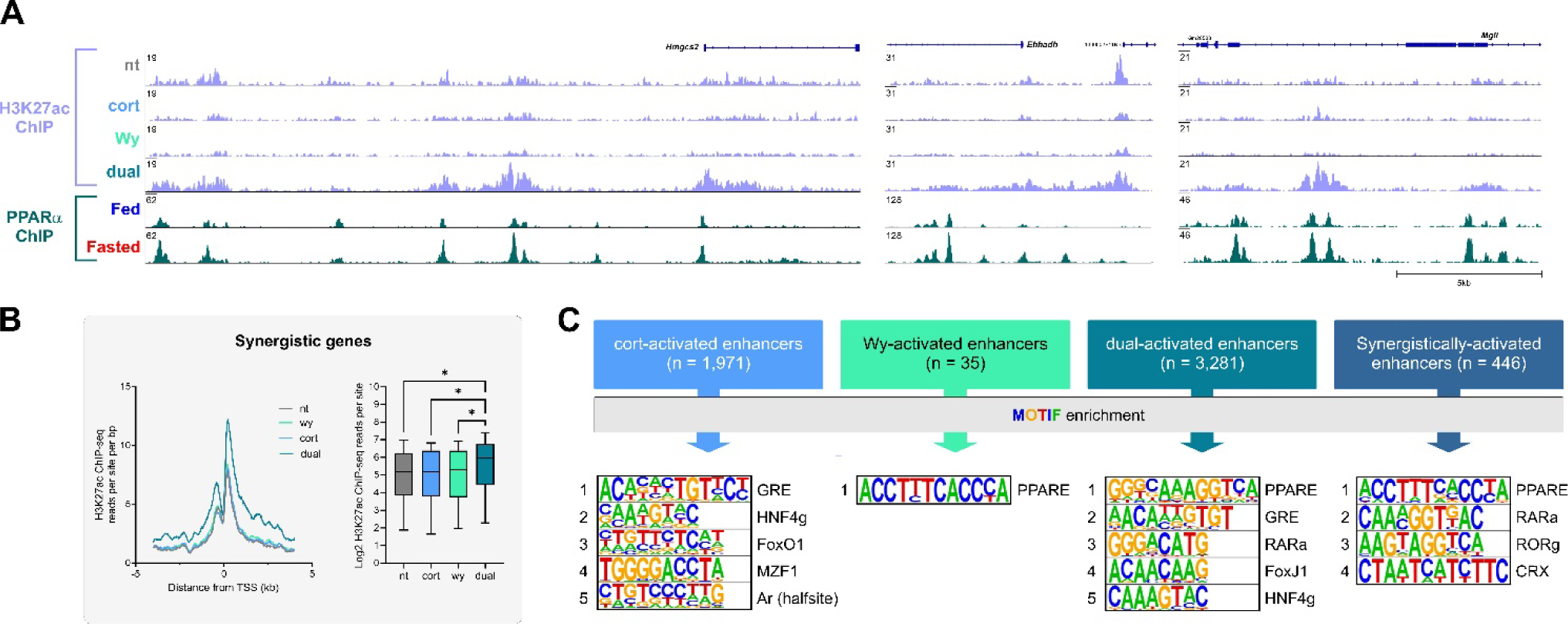
Lipid catabolism and ketogenic enhancers are synergistically-activated and are bound by PPARα. **A.** Enhancer activation patterns and PPARα binding at the loci of lipid catabolism and ketogenic genes are depicted. Enhancer activation levels, as measured by H3K27ac ChIP-seq, shows a synergistic pattern where enhancer activity level in the dual treatment markedly surpasses that of the two single treatments. Activated enhancers harbor PPARα binding sites that are overtly increased following fasting in mouse liver (PPARα ChIP-seq from [60], Fasting duration: 24 h). **B.** Quantification of H3K27ac signal around synergistically-induced genes shows a synergistic increase in H3K27ac in the dual treatment. **C.** Motif enrichment analysis of activated enhancers shows an enrichment of the GRE in corticosterone-activated enhancers, enrichment of the PPARE in Wy-activated enhancers and the enrichment of both GRE and PPARE in dual-activated enhancers. Notably, while the PPARE is enriched in synergistically-activated enhancers, the GRE is not. The complete list of enriched motifs is detailed in Table S5. - *P≤0.05, **P≤0.01, ***P≤0.001, ****P≤0.0001 by ordinary one-way ANOVA followed by Holm-Sidak post hoc analysis (B) or a two-tailed, unpaired t-test (D).

### Synergistic induction of ketogenic genes is dependent on protein synthesis and can be replaced by ectopic PPARα

In a scenario where GR-PPARα synergy is due to the GR-dependent induction of *Ppara*, one would expect that the synergistic gene induction would be somewhat delayed due to the need to synthesize the PPARα protein to sufficient amounts. Because 8 h are sufficient to accumulate PPARα (Fig. 1E), we performed a time-course experiment where hepatocytes were treated for 2, 4 or 8 h followed by gene expression quantification. We found that *Ppara* (which is a primary GR target genes as shown in Fig. 1) is quickly induced after 2 h of treatment. This is an expected kinetic behavior for primary target genes [18, 19, 63]. In contrast, the induction of synergistic genes was delayed and continually progressed with maximal induction at 4 and 8 h (Fig. 7A). Of note, this pattern does not only contrast with primary target gene induction patterns, it is also at odds with mechanisms of synergistic induction that are not dependent on transcriptional cascades as these mechanisms show peak synergistic induction at 2 h [17, 63].

**Figure 7:**
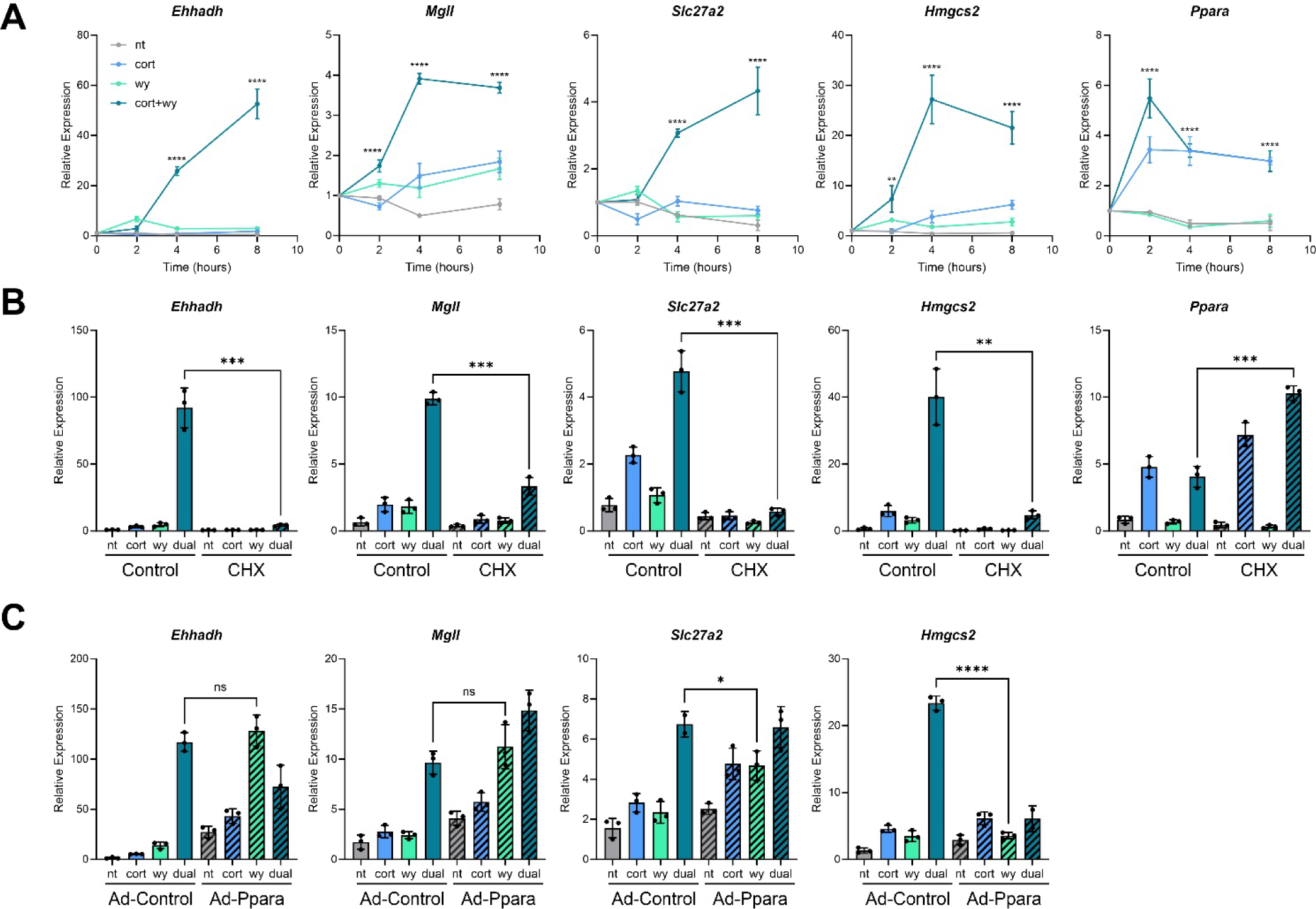
The contribution of corticosterone to synergistic gene induction is dependent on protein synthesis and is replaced by ectopic expression of PPARα. **A.** Time course experiments show corticosterone-Wy synergistic gene induction is delayed and progressively increases with time. This is in contrast to immediate and direct synergistic gene induction paradigms. Statistical tests were performed between dual-treated group and the non-treated group within each time point. **B.** Primary hepatocytes were pre-treated with a protein synthesis inhibitor (cycloheximide; CHX). Then, cells were treated with corticosterone and Wy for 8 h followed by RNA extraction and quantification. Examining gene induction patterns revealed that synergistic gene expression is dependent on protein synthesis and is abolished in the presence of CHX. **C.** Primary hepatocytes were infected with a vector expressing PPARα. In the presence of ectopically-expressed PPARα, the Wy-treated cells reached similar gene induction to that of Ad-Control cells treated with both Wy and corticosterone. This was observed for all examined genes except *Hmgcs2*. - *P≤0.05, **P≤0.01, ***P≤0.001, ****P≤0.0001 by ordinary one-way ANOVA followed by Holm-Sidak post hoc analysis (A) or a two-tailed, unpaired t-test (B, C).

The delayed pattern of synergistic gene induction, together with other evidence described above, is in line with a scenario where synergistic gene induction is due to a GR-dependent increase in PPARα protein rather that direct cooperation between GR and PPARα on enhancers. To test this, we treated hepatocytes with cycloheximide (CHX) to prevent protein synthesis. As expected, in the absence of protein synthesis the induction of *Ppara*, a primary GR target gene, is uninterrupted. However, synergistic gene induction was perturbed and the gene levels in dual-treated hepatocytes resembled those of Wy-treated hepatocytes (Fig. 7B). Thus, when PPARα synthesis is blocked, corticosterone is unable to augment the transcriptional program initiated by Wy.

Our model suggests that the major contribution of GR to synergistic gene induction is the induction of the *Ppara* gene. In this scenario, the exogenous expression of *Ppara* should be sufficient to replace the effect of corticosterone altogether. To test this, we exogenously expressed PPARα using an adenoviral vector (Ad-Ppara; Fig. S4) and treated hepatocytes with corticosterone and Wy combinations. Notably, the effect of the dual treatment in Ad-Control cells was similar to the Wy treatment in Ad-Ppara cells. In other words, the effect of corticosterone was redundant when PPARα levels are already sufficiently high (Fig. 7C). This was seen for most, but not all genes, suggesting an additional, yet unknown mechanism for GR-PPARα synergy (see Discussion for details).

In summary, in this study we found that corticosterone initiates transcriptional cascades during fasting, leading to widespread changes in enhancer activity and various gene expression programs. These cascades serve two purposes: amplifying the primary pro-gluconeogenic transcriptional response and exerting a secondary ketogenic transcriptional response. Thus, transcriptional cascades serve as a mechanism by which one hormone leads to two completely different transcriptional programs.

## Discussion

The hepatic response to fasting is heavily reliant on transcriptional and chromatin regulation with many TFs involved in it and directing expression of genes needed for fuel production. Here, we found a new manner by which fasting hormones control fuel production programs – transcriptional cascades. Glucagon and glucocorticoids each induce the expression of TF-encoding genes. We found a surprisingly high number of TF-encoding genes induced by fasting hormones and during fasting. Among the 29 induced TFs, some were previously reported to be regulated by glucagon and/or glucocorticoids (e.g. C/EBPβ [58], PPARα [56, 57, 59], KLF15 [45], FoxO1 [68, 69]) but the majority of TFs we discovered were not known to be regulated transcriptionally by these hormones. This suggests that transcriptional cascades are a major part of the transcriptional response to fasting. Indeed, we showed two transcriptional cascades that are profoundly involved in directing fuel production gene programs following their induction.

We revealed that a major role of C/EBPβ in hepatocytes is to amplify gene induction initiated by glucagon and corticosterone, thereby amplifying a gluconeogenic gene program and augmenting gluconeogenesis. In addition, C/EBPβ amplifies other hepatic gene programs driven by glucagon and glucocorticoids such as bile acid metabolism, cholesterol transport and the acute phase response. As such, these findings place C/EBPβ as an amplifier of hormone-dependent gene induction. It is conceivable that this amplification is brought about by cooperation of C/EBPβ with hormone-activated TFs: CREB and GR. Indeed, such cooperation was reported where C/EBPβ was found to assist the loading of GR to enhancers. GR binding was increased in the presence of C/EBPβ at a specific set of enhancers co-bound by both C/EBPβ and GR. Assisted loading was associated with a C/EBPβ-dependent increase in enhancer accessibility [70].

In contrast to C/EBPβ that amplifies the primary transcriptional program, PPARα exerts a secondary gene program aimed at inducing FAO and ketogenic genes. While PPARα activity is absolutely dependent on ligand activation, we found that glucocorticoids presence is a prerequisite to its function. A dual treatment of corticosterone and Wy led to an overt synergistic response with 80 genes synergistically induced. The synergy between the glucocorticoid-GR axis and PPARα has been widely described [18, 59, 71–73]. We show here a far-reaching effect of the dual corticosterone-Wy treatment on hepatic enhancer landscape with 446 enhancers synergistically activated. Importantly, we provide several evidence that the induction of *Ppara* by glucocorticoids greatly contributes to this synergy: (a) synergistic gene induction is delayed, fitting with the need to synthesize protein before synergy can take place. (b) synergistically-activated enhancers are enriched for PPARE and not GRE, arguing against cooperative binding of PPARα and GR. (c) inhibition of protein synthesis negates synergistic gene induction. (d) ectopic expression of PPARα makes corticosterone redundant, suggesting the main role of corticosterone is to increase PPARα levels.

While the GR-PPARα transcriptional cascade is a major determinant in synergistic gene induction, we assume additional mechanisms are at play. This is because not all synergistic gene induction was fully recapitulated by ectopic PPARα expression. It is tempting to speculate that assisted loading between GR and PPARα take place at some enhancers, leading to synergistic gene induction, as we and other showed in other synergistic paradigms [17, 18, 63, 74]. Unfortunately, we were unable to profile PPARα via ChIP-seq in primary hepatocytes and therefore this possible mechanism remains a hypothesis for the moment.

Taken together, our findings place transcriptional cascades as a key mechanism driving fuel production during fasting. transcriptional cascades serve both to amplify a primary response as well as to generate a delayed secondary transcriptional response. Our observations portray transcriptional cascades as an efficient mechanism to respond to endocrine and other extracellular signals. Transcriptional cascades provide both a multi-layered and temporally-organized response to signals, thus tailoring an adaptive response to a changing environment.

## Materials and methods

### Reagents

Glucagon 100 nM (Ray biotech, cat# 228-10549-1), corticosterone 1 µM (Sigma-Aldrich, cat# 50-22-6), Wy14643 10 µM (TCI chemicals, cat# C1323), RU486 2 µM (Sigma, cat# M8046), CHX 1 µg/ml (Sigma, cat# 01810).

### Primary mouse hepatocytes

primary hepatocytes were isolated from male, 8-10 weeks-old mice (strain C57BL/6JOlaHsd). Isolation and plating of primary mouse hepatocytes was performed as detailed in our published protocol with no modifications [75]. Three hours after plating, media was changed to Williams E media (ThermoFisher Scientific, cat# 12551032). All hormone treatments were performed 18 h after plating in Williams E media except for adenovirus infection experiments, detailed below. All animal procedures are compatible with the standards for care and use of laboratory animals. The research has been approved by the Hebrew University of Jerusalem Institutional Animal Care and Use Committee (IACUC). The Hebrew University of Jerusalem is accredited by the NIH and by AAALAC to perform experiments on laboratory animals (NIH approval number: OPRR-A01-5011).

### RNA preparation, reverse transcription and quantitative PCR (qPCR)

Total RNA was isolated from primary mouse hepatocytes using NucleoSpin kit (Macherey-Nagel cat# 740955.25) according to the manufacturer’s protocol. For qPCR, 1 μg of total RNA was reverse transcribed to generate cDNA (Quantabio cat# 76047-074). qPCR was performed using a C1000 Touch thermal cycler CFX96 or CFX Opus 384 thermal cycler instruments (Bio-Rad) using SYBR Green (Quantabio cat# 101414-276). Gene values were normalized with a house keeping gene (*Rpl13*). Primers were designed to amplify nascent transcripts (i.e., the amplified region span exon-intron junctions) as a proxy for transcription and in order to avoid confounding post-transcriptional events. This was done for all primers except for Cebpb which is intronless and Rpl13 which was used only for normalization levels. The sequences of primers used in this study are:

*Rpl13* - Fwd: AGCCTACCAGAAAGTTTGCTTAC, Rev: GCTTCTTCTTCCGATAGTGCATC
*Cebpb* - Fwd: ATCCGGATCAAACGTGGCTG, Rev: GGCCCGGCTGACAGTTAC
*Ppara* - Fwd: GAGGCTTACCCAGAGTCCAC, Rev: ATCTTGCAGCTCCGATCACA
*Ehhadh* - Fwd: AAACACTGTCCATGTTGGGC, Rev: CACCGCACTGTCGTTTTGTTT
*Hmgcs2* - Fwd: CACTCTACTCTGGATTTGGCAGT, Rev: CCACTCGGGTAGACTGCAATG
*Mgll* - Fwd: CTAAGCAACACGGCAACGTC, Rev: CTGTGGAGGACGTGATAGGC
*Slc27a2* - Fwd: GGTCCGTGACGCAAATGGAT, Rev: GCACGGTTAATTGCGCAGAG

### Chromatin immunoprecipitation (ChIP)

PMH were treated with treatment combinations for 8 h. Then, cells were cross-linked with 1% formaldehyde (Electron Microscopy Sciences, cat# 15714) for 10 min at room temperature and quenched with 0.125M glycine. Crosslinked samples were washed in phosphate buffered saline (PBS), resuspended in ChIP lysis buffer (0.5% SDS, 10mM EDTA, 50mM Tris-HCl pH8) and sonicated (Bioruptor Plus, Diagenode) to release 100-1000 bp fragments. Antibody (4 μl per sample) against H3K27ac (Active Motif, cat# 39133) was conjugated to magnetic beads (Sera-Mag, Merck, cat# GE17152104010150) for 2 h at 4°C. Chromatin was immunoprecipitated with antibody-bead conjugates for 16 h at 4°C. Immunocomplexes were washed sequentially with the following buffers: low-salt buffer (0.01% SDS, 1% Triton x-100, 2 mM EDTA, 20mM Tris-HCl pH8, 150mM NaCl), high salt buffer (0.01% SDS, 1% Triton x-100, 2mM EDTA, 20mM Tris-HCl pH8, 500mM NaCl), low salt buffer and TE buffer (10mMTris-HCl, 1mM EDTA pH8). Chromatin was de-proteinized with proteinase K (Hy Labs, cat# EPR9016) for 2 h at 55°C and de-crosslinked for 12 h at 65°C. DNA was subsequently phenol-chloroform purified and ethanol precipitated.

### RNA-seq and ChIP-seq

For quality control of RNA yield and library synthesis products, the RNA ScreenTape and D1000 ScreenTape kits (both from Agilent Technologies), Qubit RNA HS Assay kit, and Qubit DNA HS Assay kit (both from Invitrogen) were used for each specific step. mRNA libraries were prepared from 1 µg RNA using the KAPA Stranded mRNA-Seq Kit, with mRNA Capture Beads (KAPA biosystems, cat# KK8421). ChIP DNA libraries were prepared using the KAPA HyperPrep Kit (KAPA biosystems, cat# KR0961). The multiplex sample pool (1.6 pM including PhiX 1%) was loaded on NextSeq 500/550 High Output v2 kit (75 cycles) cartridge, and loaded onto the NextSeq 500 System (Illumina), with 75 cycles and single-read sequencing conditions.

### Western blot

To prepare whole cell lysates, RIPA (50 mM tris-Cl, 150 mM NaCl, 1% triton, 0.5% sodium deoxycholate, 0.1% SDS) was added directly on adherent cells followed by scraping and centrifugation (4°C, 2000 rpm, 10 min.). Protein samples were loaded on SurePAGE™ Bis-Tris, 10×8, 4-20% polyacrylamide SDS gels (GeneScript, M00657). Proteins were transferred (Trans Blot Turbo, Bio-Rad; cat# 1704158) to a nitrocellulose membrane (Trans-Blot Turbo Transfer Pack, Bio-Rad; cat# 1704158), blocked for 1 hour with 5% low-fat milk, and incubated for 16 h with primary antibody (PPARα - Santa Cruz Biotechnology cat# sc-398394, H3 - Cell Signaling Technologies cat# 14269, CEBPβ - Santa Cruz Biotechnology cat# sc-150) diluted 1:1000 in solution (tris-buffered saline 0.5% Tween, 5% bovine serum albumin, 0.05% sodium azide). Membranes were incubated with secondary peroxidase AffiniPure goat anti-rabbit immunoglobulin G (1:10,000, Jackson Laboratory, cat# 111-035-144) or anti-mouse (1:10,000, Jackson Laboratory; cat# 115-035-146) for 1 h, followed by washes and a 1-minute incubation with western blotting detection reagent (Cytiva Amersham ECL prime, cat# RPN2232). Imaging and quantification were done with ChemiDoc (Bio-Rad).

### Adenovirus infection

Three hours after plating, PMH were infected with adenovirus: Ad-ShCebpb (Vector Biolabs, cat# shADV-255244, concentration: 25000 PFU/µl) or Ad-Ppara (Vector Biolabs, cat# ADV-269120, concentration: 10000 PFU/µl). After 12 h (Ad-shCebpb) or 16 h (Ad-Ppara), hormones were added for downstream experiments.

### Hepatic glucose production

Twenty-four hours after plating, primary hepatocytes (plated in 12-well plates; 2.5 × 10^5^ cells per well) were incubated for 9 hours with Dulbecco modified Eagle medium lacking glucose, pyruvate, glutamine, and phenol red (Gibco, cat# A14430-01). Then cells were washed with PBS and treated for 20 hours with a medium containing a combination of glucagon and corticosterone together with L-Glutamine (20 mM, Sigma, cat# 59202C): After 20 hours, 40 µL of medium was sampled, and glucose was measured using the glucose oxidase colorimetric method according to the manufacturer’s instructions (Sigma, cat# GAGO20).

### Sequencing data analyses

Fastq files were mapped to the mm10 mouse genome assembly using Bowtie2 [76] with default parameters. Tag directories were made using the makeTagDirectory option in HOMER [77]. H3K27ac peaks were called using MACS2 (narrowPeak option) [78]. All site overlaps were performed by MergePeaks option in HOMER. Selected gene loci were visualized by the integrated genome browser (IGV) [79].

### Differential gene expression

Differential gene expression was evaluated by DEseq2 [80] via the HOMER suite under default parameters. Genes were determined as differentially expressed between two conditions if they pass these cutoffs: fold change ≥ or ≤ 1.5, adjusted p value ≤ 0.05.

### k-means clustering

All C/EBPβ-regulated genes (both upregulated and downregulated) were included in the clustering analysis fold change ≥ 1.5, adj. p value ≤ 0.05). The normalized tag counts of each gene were used for the analyses. Morpheus (https://software.broadinstitute.org/morpheus) was used to cluster genes under these parameters: k = 8; metric - one minus Pearson correlation; maximum iterations – 1000. Blue - minimum value of the gene; Red – maximum value of each gene (minimum and maximum values of each gene are set independently to other genes).

### H3K27ac ChIP-seq analyses

Peak-calling was performed by MACS2, sites common to at least two replicates were merged and ENCODE blacklisted sites were omitted. Differential enhancer activity (DEseq2 fold change ≥ 1.5, adj. p value ≤ 0.05) was measured in all conditions as compared to the non-treated control.

### De novo motif enrichment analysis

To unbiasedly detect enriched motifs, we performed a de novo motif enrichment analysis using the findMotifsGenome option in HOMER (parameter -size given). The entire enhancer landscape (all H3K27ac sites appearing in at least two replicates across all conditions) was used as background to account for possible sequence bias. Using the entire enhancer landscape as background ensures that prevalent motifs appearing across liver enhancers will not be falsely detected as specifically enriched in the examined subset of enhancers. All motifs with p value ≤ 1^−10^ are presented.

### Aggregate plots and box plots

Tag density of H3K27ac, C/EBPβ and PPARα signal around enhancers or transcription start sites (TSS) were analyzed using the HOMER suite. In aggregate plots, the tag count (averaged across all sites) per site per bp was calculated using the HOMER suite (annotatePeaks, option -size 8000 -hist 10). In box plots, tag count +/− 200 bp around the site center (averaged across all sites) was calculated using the HOMER suite: annotatePeaks, option -size 400 -noann. In box plots for H3K27ac, which is a broader signal, tag count +/− 500 bp around the site center (averaged across all sites) was calculated using the HOMER suite: annotatePeaks, option -size 1000 -noann. In both aggregate plots and box plots, the data is an average of all three replicates. In all box plots, the 10-90 percentiles are plotted.

### GO terms enrichment analysis

performed by GeneAnalytics [81]

### Analyses of published data

In addition to data generated in this study, we used data that was previously published by us [17–19] and others [60].

### Statistical analyses

All conditions in all of the described experiments were performed in three biological replicates. Error bars represent standard deviation of biological replicates. In pairwise comparisons, statistical significance was determined by a two-tailed, unpaired t-test. In comparisons of three or more groups, ordinary one-way ANOVA was performed with post-hoc analyses made via Holm-Sidak tests as specified in the figure legends. In RNA-seq and ChIP-seq experiments, DESeq2 was used to determine statistical significance. *P≤0.05, **P≤0.01, ***P≤0.001, ****P≤0.0001, ^ns^ P> 0.05. Further details about statistical analyses are described in figure legends.

## Acknowledgments

We would like to thank Dr. Abed Nasereddin and Dr. Idit Shiff from The Genomic Applications Lab, Hebrew University of Jerusalem, for their invaluable help, support and expertise in high throughput sequencing.

## Funding

The Israel Science Foundation (ISF, grants numbers 1469/19, 3533/19); European Research Council (ERC-StG grant number 947907).

## Author contributions

Contribution categories were adopted from CRediT (Contributor Roles Taxonomy)

D. G. was involved in Conceptualization, Methodology, Software, Validation, Formal analysis, Investigation, Writing, and Visualization

N. B. was involved in Investigation

M. C-N was involved in Conceptualization, Methodology, Software, Formal analysis, and Investigation

I. G was involved in Conceptualization, Methodology, Software, Validation, Formal analysis, Investigation, Resources, Writing, Visualization, Supervision, Project administration, and Funding acquisition.

## Conflict of interests

The authors declare that they have no conflict of interest

## Data availability

All RNA-seq and ChIP-seq data have been deposited in the Gene Expression Omnibus (GEO; https://www.ncbi.nlm.nih.gov/geo/) under accession number GSE252320.

## Notes

### Competing Interest Statement

The authors have declared no competing interest.

