## Supplementary figures and images for "Transcriptional cascades during fasting amplify gluconeogenesis and instigate a secondary wave of ketogenic gene transcription"

### Figure S1

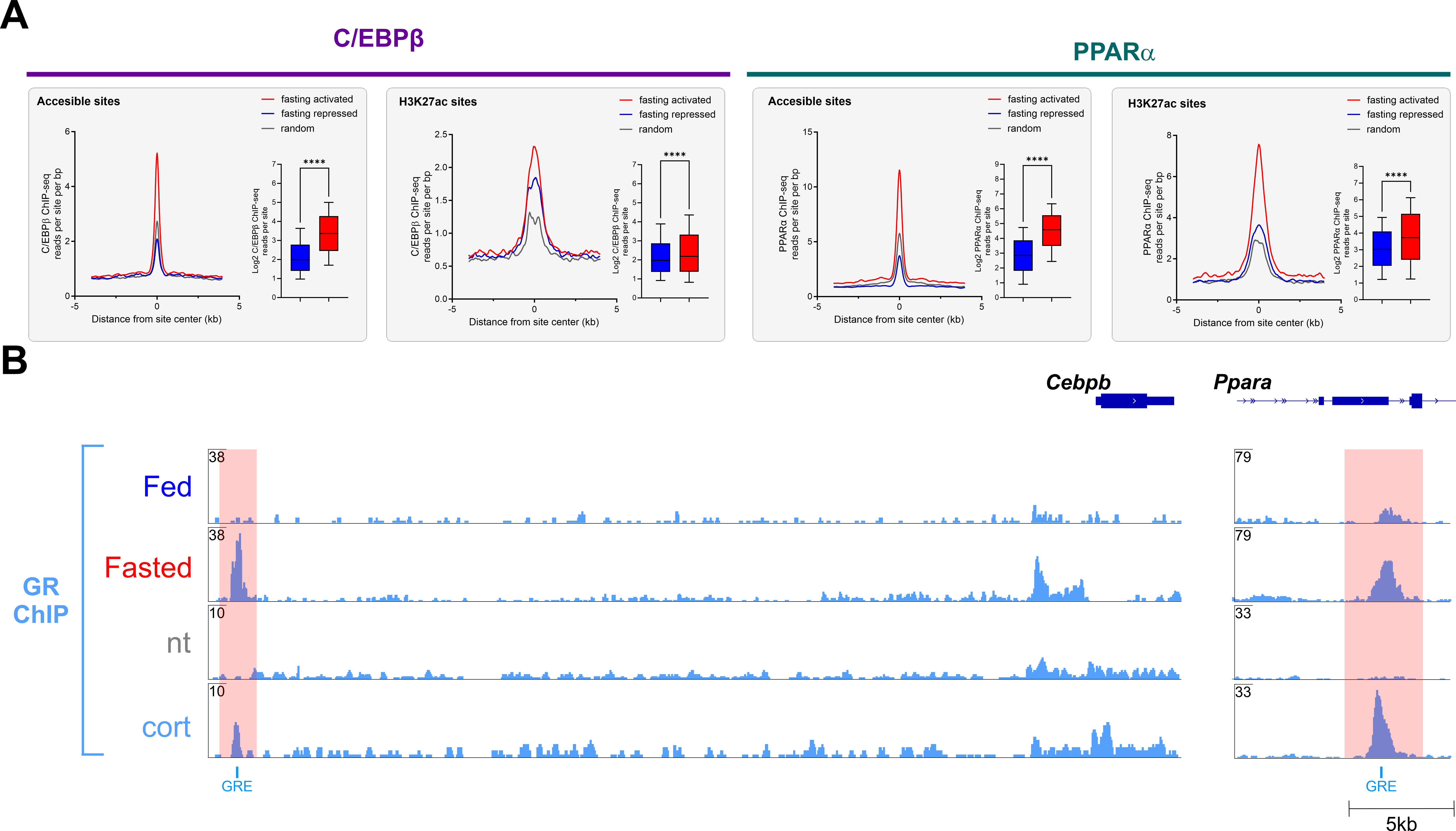

### Figure S2

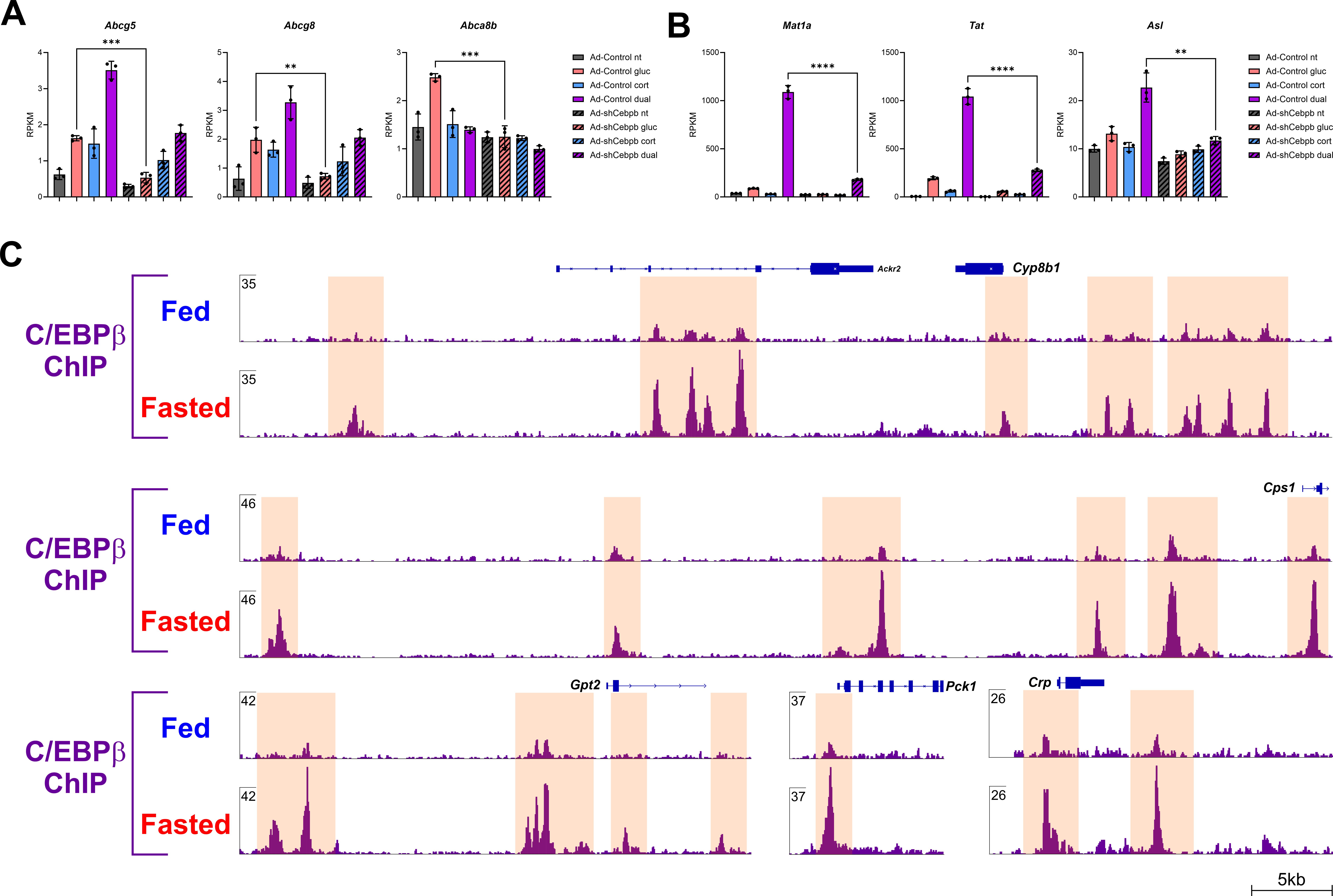

### Figure S3

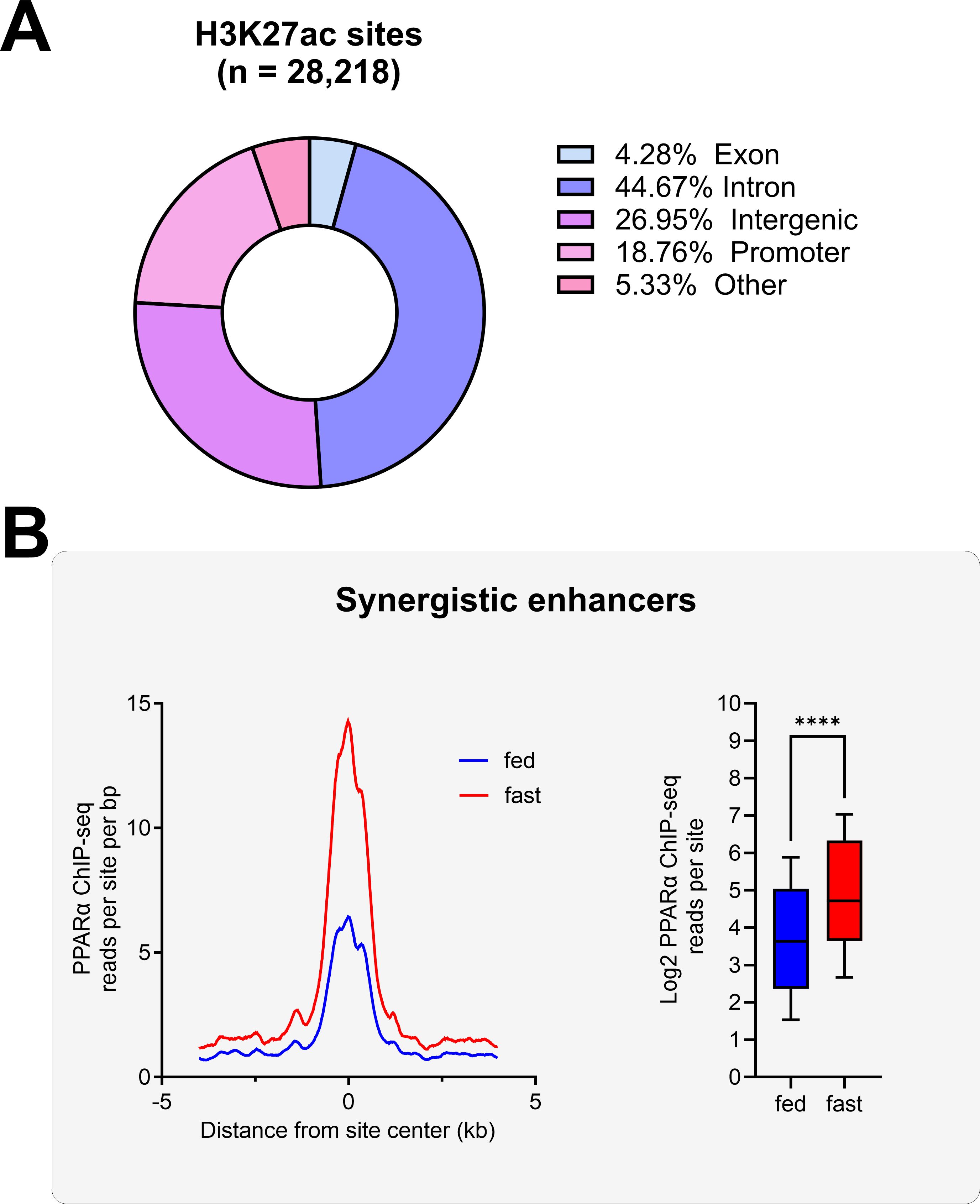

### Figure S4

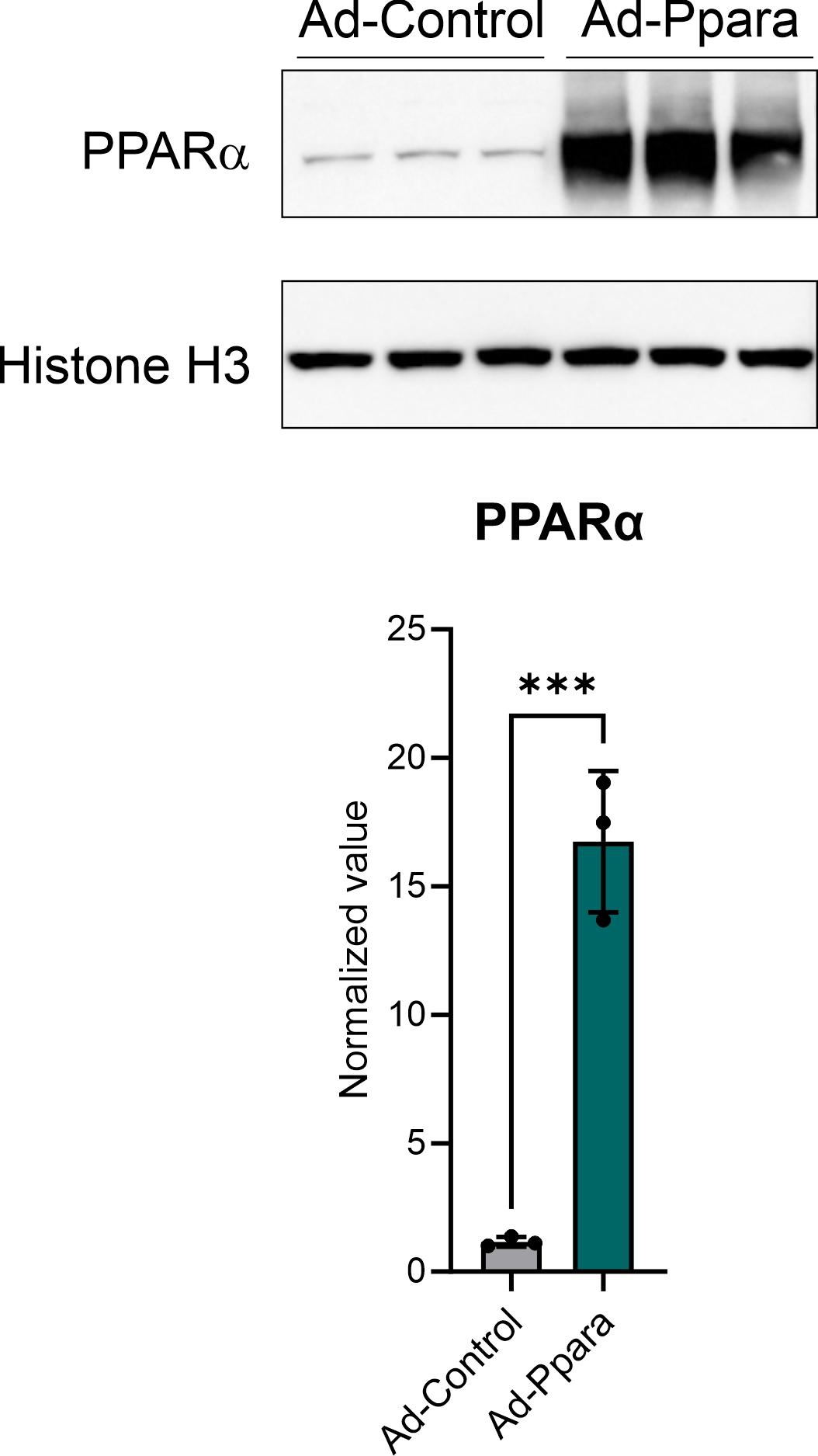
